# Identification of changing ribosome protein compositions using cryo-EM and mass spectrometry

**DOI:** 10.1101/271833

**Authors:** Ming Sun, Parimal Samir, Bingxin Shen, Wen Li, Christopher M. Browne, Rahul, Joachim Frank, Andrew J. Link

**Author notes:** Equal contributions (Ming Sun, Parimal Samir, Bingxin Shen). Corresponding authors Andrew J. Link Dept. of Pathology, Microbiology and Immunology Vanderbilt University School of Medicine 1161 21^st^ Ave South Nashville, TN 37232, Joachim Frank Department of Biochemistry and Molecular Biophysics Columbia University 650 W. 168^th^ Street New York, NY 10032.

## Abstract

The regulatory role of the ribosome in gene expression has come into sharper focus. It has been proposed that ribosomes are dynamic complexes capable of changing their protein composition in response to enviromental stimuli. We applied both cryo-EM and mass spectrometry to identify such changes in S. cerevisiae 80S ribosomes. Cryo-EM shows a fraction (17%) of the ribosome population in yeast growing in glucose lack the ribosomal proteins RPL10 (ul16) and RPS1A/B (eS1). Unexpectedly, this fraction rapidly increases to 34% after the yeast are switched to growth in glycerol. Using quantitative mass spectrometry, we found that the paralog yeast ribosomal proteins RPL8A (eL8A) and RPL8B (eL8B) change their relative proportions in the 80S ribosome when yeast are switched from growth in glucose to glycerol. Using yeast genetics and polysome profiling, we show that yeast ribosomes containing either RPL8A or RPL8B are not functionally interchangeable. Our combined cryo-EM and quantitative proteomic data support the hypothesis that ribosomes are dynamic complexes that alter their composition and functional activity in response to changes in growth or environmental conditions.

## Highlights

- A fraction of ribosomes in yeast cells growing in glucose lacks two essential proteins and appears inactive.
- This inactive fraction doubles when the cells are switched from glucose to glycerol as the carbon source.
- The switch in carbon source induces compositional changes in ribosomal protein paralogs in the ribosomes and functional changes based on polysome profiles.
- RPL8A and RPLB change their relative stoichiometry in the ribosome when yeast cells are grown in different carbon sources.

## Introduction

Gene expression can be regulated at multiple levels, including transcription and translation. Translation is the process by which the information encoded in an mRNA is used to synthesize polypeptides. Composed of a small 40S and large 60S subunit, the eukaryotic 80S ribosome catalyzes the decoding of mRNAs and formation of peptide bonds. Translational control is a major mechanism modulating eukaryotic gene expression (**Carpenter** *et al.*, 2014; **Costa-Mattioli** *et al.*, 2009; **Hinnebusch**, 2015; **Holcik** and Sonenberg, 2005; **Kong** and Lasko, 2012). Numerous studies have shown that a large number of translation factors, posttranslational protein modifications, and RNA accessory factors play active roles in translational control (**Dever** and Green, 2012; **Hinnebusch**, 2015; **Kapp** and Lorsch, 2004; **Sonenberg** and Hinnebusch, 2009). Initially, the ribosomes themselves were considered only passive players in this process (**Mauro** and Edelman, 2002).

In recent years, the idea that the ribosomes can function as transcript-specific posttranscriptional regulatory elements has been formulated (**Komili** *et al.*, 2007; **Kondrashov** *et al.*, 2011; **McIntosh** and Warner, 2007; **Ruggero** and Pandolfi, 2003; **Warner**, 2015; **Warner** and McIntosh, 2009; **Xue** *et al.*, 2015). In the depot model, ribosomes act as reservoirs of regulatory molecules, which are released in response to specific cellular cues (**Mazumder** *et al.*, 2003; **Ray** *et al.*, 2007; **Zhou** *et al.*, 2015). An example of this mode of action is the role of the human RPL13A ribosomal protein (r-protein) in translational control during the inflammatory response (**Kapasi** *et al.*, 2007; **Mazumder** *et al.*, 2003). A second model proposes a more direct role of ribosomes in translational control (**Mauro** and Edelman, 2002; **Mauro** and Edelman, 2007). This model derives from studies that reveal ribosomes are not the static, uniform structures described in textbooks. Comparison of ribosomes from rat skeletal muscle and liver using 2-D gel electrophoresis revealed differences in ribosomal composition between the two tissues (**Sherton** and Wool, 1974). In the amoeba *Dictyostelium discoideum* ribosomes from spores and vegetative cells differ both in protein composition and posttranslational modifications (**Ramagopal** and Ennis, 1981). In the *S. cerevisiae* genome, the 79 r-proteins in the 80S ribosome are encoded by 138 genes (**Nakao** *et al.*, 2004; **Table S1**). These include 57 duplicate, paralogous ribosomal gene pairs (**McIntosh** and Warner, 2007; **Warner**, 1999; **Table S1**), raising the possibility that individual paralogs have distinct finctions. Deletion analysis of *S. cerevisiae* paralogous pairs showed that the genes are not functionally equivalent (**Breslow** *et al.*, 2008; **Giaever** *et al.*, 2002; **Komili** *et al.*, 2007). Using mass spectrometry, differences were found in the stoichiometry of core r-proteins in yeast grown under different growth conditions (**Slavov** *et al.*, 2015). In higher eukaryotic organisms, previous studies showed specific r-proteins are required for the translation of selected mRNA transcripts (**Kondrashov** *et al.*, 2011; **Lee** *et al.*, 2013). Post-translational modifications, especially phosphorylation, and variations in rRNA composition add further levels of complexity to the different ribosome complexes in cells and tissues (**Martin** *et al.*, 2014; **Ramagopal**, 1992).

The evidence of heterogeneity in the ribosome’s protein and rRNA composition has generated speculation as to its functional consequences. Mauro and Edelman proposed that ribosomal subunits differing in protein or rRNA composition bind translation factors or specific mRNAs with different affinities, thereby selectively changing the rates of polypeptide synthesis of particular mRNAs (**Mauro** and Edelman, 2002; **Mauro** and Edelman, 2007). In their model, different or specialized ribosomes in a heterogeneous ribosomal population in the cell function as regulatory elements or ‘filters’ by selectively binding and translating specific mRNAs. In support of this model, Bauer *et al.* showed that “specialized” ribosomes with specific r-proteins encoded by paralogous genes preferentially translate particular reporter mRNAs (**Bauer** *et al.*, 2013).

Cells growing under one growth condition require a specific proteome, and the cells growing in a second condition require a different proteome. A number of cellular mechanisms including transcription, RNA processing, translation, and protein degradation are known to regulate gene expression and ultimately the cell’s proteome when growth conditions change. In this study, we hypothesized that the cell’s ribosomes also change their protein composition in response to changing growth conditions as a mechanism to regulate expression. To test this hypothesis, we used a combined strategy of single-particle cryo-electron microscopy (cryo-EM) and quantitative mass spectrometry-based proteomics to detect and identify specific r-proteins that change their stoichiometry in the 80S ribosome upon change of growth condition. Cryo-EM is a method for imaging molecules in their native, hydrated state at cryogenic temperature by transmission electron microscopy (**Frank**, 2006). When applied to the ribosome, cryo-EM can be used to reconstruct the ribosome at molecular resolution in three dimensions. Moreover, the technique allows ribosomes to be grouped into different classes, or subpopulations, based upon structural similarities and differences. Reconstructions of the individual classes can be compared with a high-resolution reference structure of the intact 80S ribosome to identify subpopulations lacking densities for r-proteins. In this way, cryo-EM can be used to detect and identify ribosomes with sub-stoichiometric r-protein compositions. Quantitative proteomics using tandem mass spectrometry analysis of protease-digested ribosomes coupled with isotopic labeling can be used to both identify and quantify the relative abundances of specific proteins in cellular or subcellular protein fractions (**Ross** et al. MCP 3, 1154, 2004; **Yates** *et al.*, 2009). Thus, cryo-EM and quantitative proteomics should be able complement each other’s strengths in a comprehensive approach to identify changes in 80S ribosomal protein compositions.

Using such a combined strategy, we profiled the 80S ribosome composition in yeast cells growing in glucose or glycerol as a carbon source. Abruptly shifting from fermentable glucose to the non-fermentable glycerol carbon source induces a diverse array of changes in *S. cerevisiae* gene expression (**Kuhn** et al MCB, 21, 916, 2001; **Samir** *et al.*, 2015). In this study, ribosomes from yeast cells growing in glucose and then glycerol were isolated using sucrose gradients and split into two pools for cyro-EM and quantitative proteomics. Cryo-EM identified a significant fraction of 80S ribosomes from glucose growth lacking the essential r-protein RPL10 or both RPL10 and RPS1A/B. This fraction rapidly increased upon switching to growth in glycerol. Importantly, these classes of ribosomes also lacked tRNAs, as well as any conformational variability known to be associated with active translation. Using proteomic analysis, we identified 75 of the expected 79 r-proteins in the 80S complexes. This set included 55 of the 57 paralogous r-protein gene pairs. Proteomic analysis revealed that the r-protein paralogs RPL8A and RPL8B change their relative stoichiometry in the population of ribosomes when yeast cells are shifted from glucose to glycerol as a carbon source. Phenotypic analysis and polysome profiling using *rpl8a*Δ and *rpl8b*Δ null homozygotes show that the functions of RPS8A and RPS8B are not interchangeable.

## Results

We hypothesized that specific r-proteins important for the specialization of ribosome activity will show differential abundances in response to environmental stimuli. In pilot studies, we first used quantitative proteomics to identify and quantify changes in individual r-proteins abundances from yeast whole-cell protein extracts after shifts in both carbon source and temperature (**Samir** *et al.*, 2015). The analysis showed most of the 40S and 60S r-proteins were down-regulated in response to the two environmental stimuli (**Fig. 1**). However, selected r-proteins showed significantly larger decreases in abundance (**Samir** *et al.*, 2015). This result suggested that these r-proteins were being differentially expressed compared to the majority of the r-proteins, consistent with the ribosome filter hypothesis (Mauro and Edelman, 2002).

**Fig. 1.**
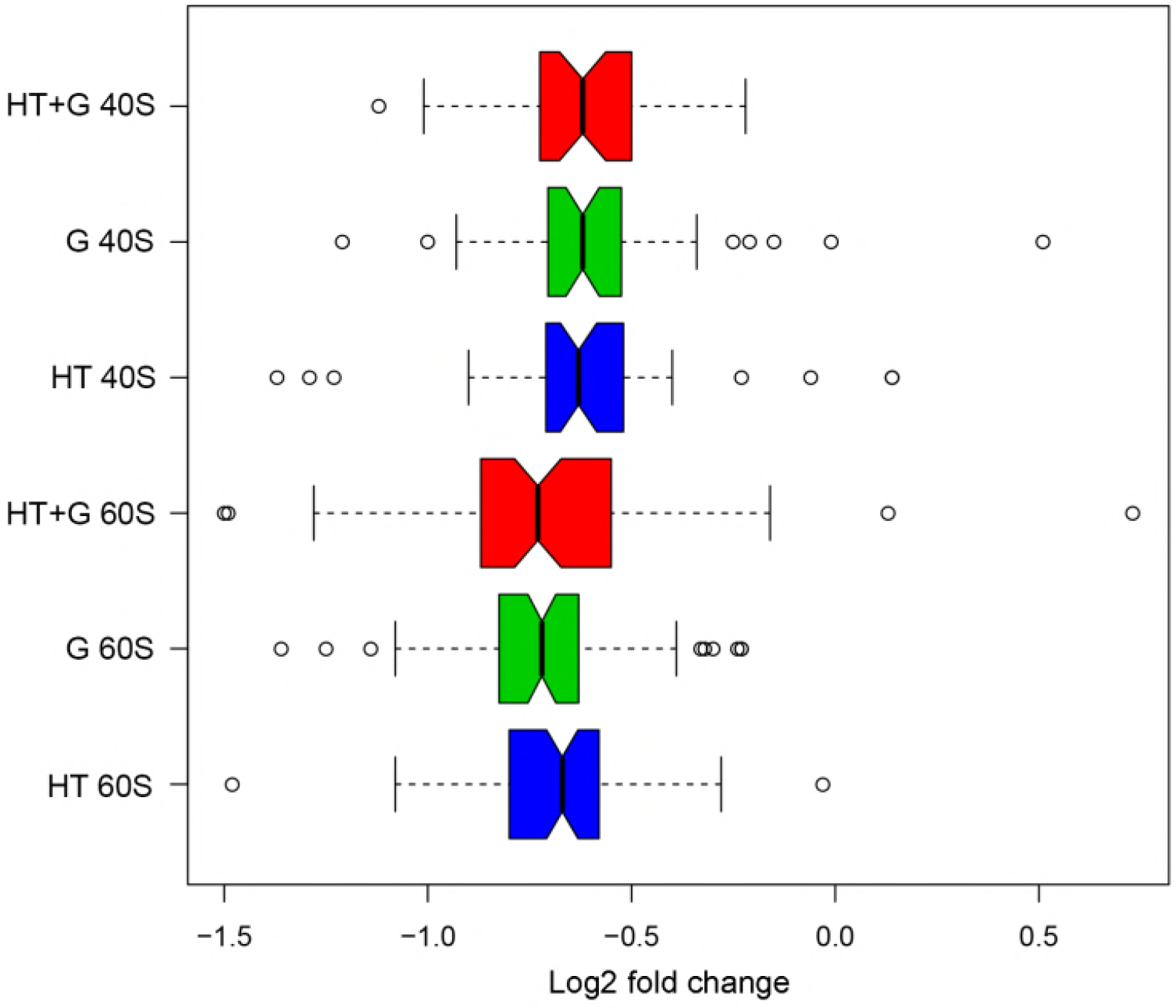
The log_2_ fold-changes in the distributions of 40S and 60S subunit protein abundances caused by changes in growth conditions. Quantitative proteomics was used to identify and quantify changes in individual r-proteins abundances from *S. cerevisiae* whole-cell protein extracts after shifts in carbon source and temperature (**Samir** *et al.*, 2015). Yeast was grown at standard conditions (glucose, 30°C) and changed to three different environmental growth conditions or stimuli: high temperature 39°C (HT, blue); glycerol (G, green), and high temperature and glycerol combined (HT+G, red). A custom Python script *CompZilla.py* was used to analyze changes in the r-proteins in the quantitative proteomic data. Open circles represent r-protein outliers in the log_2_ fold-changes in their protein abundance.

In order to directly measure the changes in the protein composition of 80S ribosomes purified from yeast, we focused on comparing yeast growing in rich medium with two different carbon sources, glucose and glycerol (**Samir** *et al.*, 2015). From yeast cells cultured in glucose or glycerol, we isolated ribosomes using discontinuous sucrose gradient centrifugation to use for both cryo-EM and quantitative proteomic analysis (**Samir** *et al.*, 2015).

### Cryo-EM studies of *S. cerevisiae* ribosomes at multiple time points

To study the changes in the structure and composition of 80S ribosomes over time after changing from glucose to glycerol, we first performed cryo-EM experiments on 80S ribosomes harvested from ribosomal samples at different time points. An aliquot of cells growing in glucose was collected (T=0 min) and the cells were shifted to growth in glycerol. We then harvested cells at 30, 120, 240, 450 and 1440 min after shifting to glycerol and purified ribosomes using sucrose gradients. The cryo-EM study proceeded in two stages. In the initial stage, the method of classification and quantification of ribosome subpopulations was developed and validated with a dataset collected on a 200-kV instrument using a CCD camera (**Shen**, B, Sun, M., Li, W., Samir, P., Link, A. and Frank, J., in preparation). In the present full-scale study, a much larger dataset was collected on a 300-kV instrument using a direct electron detection camera.

***Initial cryo-EM study***. Initial cryo-EM experiments were performed on the purified yeast 80S ribosomes after the shift from glucose to glycerol, using an FEI Tecnai F20 electron microscope (FEI, Portland, OR) at 200 kV, equipped with a 4k × 4k CCD camera (Gatan, Pleasanton, CA). A total of ~121,000 particles were selected from 2,661 micrographs from the six time-resolved sample sets. Hierarchical classification was performed using the *RELION* program (**Scheres**, 2012), in conjunction with a quantitative analysis of convergence (**Chen** *et al.*, 2014). To quantitatively analyze the results, we used a pooled classification scheme that tracks the shifts of sub-populations and ensures the maximum number of particles for the reconstruction of each ribosome class (**Shen**, B, Sun, M., Li, W., Samir, P., Link, A. and Frank, J., in preparation). The majority of the cryo-EM ribosome images at T=0 were of complete 80S ribosomes, bound with P/E-tRNA, A/P-P/E-tRNAs or P-E-tRNAs, strongly suggesting they were biologically functional. However, a fraction (~20%) lacked the density for r-proteins RPS1A/B (eS1) and RPL10 (uL16) and showed no tRNA. Unexpectedly, 30 min after shifting from glucose to glycerol, the fraction of 80S ribosomes lacking these r-proteins increased considerably (to ~32%) and remained at this level.

***Full-scale cryo-EM study***.Prompted by the observation of the change in r-protein composition of the 80S ribosomes, we used an FEI Polara TF30 electron microscope operating at 300kV, equipped with GATAN K2 Summit direct detector camera (Gatan, Pleasanton, CA), to collect a larger dataset in an attempt to improve the resolution. A total number of ~291,000 particles were selected from 6,576 micrographs from the six sample sets (i.e., glucose at T=0 and five different time points after the switch to growth in glycerol). The resulting six cryo-EM data sets were pooled together for further classification and refinement, following the protocol established in the initial study (**Shen**, B, Sun, M., Li, W., Samir, P., Link, A. and Frank, J., in preparation; for an overview of the data processing procedure, see **Fig. S1**). The first tier in the hierarchical unsupervised classification revealed two major ribosomal populations, complete 80S ribosomes and incomplete 80S ribosomes lacking densities for r-proteins RPS1A/B (eS1) and/or RPL10 (uL16).

Since reconstructions from both populations showed fragmented densities, we performed a second run of exhaustive unsupervised classification. For the population of complete 80S ribosomes, we identified five major subpopulations differing in conformation and occupancy by tRNAs (Fig. S1 **and Table S2**). We also identified two types of conformational changes, inter-subunit rotation and small subunit head swivel, which are known to accompany mRNA-tRNA translocation. The findings of tRNAs and their associated conformational changes is evidence of actively translating ribosomes (**Frank** and Gonzalez, 2010). For brevity, 80S ribosomes with and without evidence of inter-subunit rotation will be termed “rotated” and “non-rotated,” respectively. Three (i-iii) of the five subpopulations showed rotated 80S ribosomes bound with hybrid P/E tRNA (denoted as rt80-P/E), differing in the extent of their 40S subunit head swivel. One (iv) showed non-rotated 80S ribosomes bound with classically configured P-and E-tRNAs (denoted as nrt80S-P/E); and one (v) showed non-rotated 80S without tRNAs (denoted as nrt80S-empty). The resolutions of the density maps reconstructed from these subclasses ranged from 5.6Å to 7.6Å (here and henceforth, all resolutions were measured by FSC=0.143, following the ‘gold-standard’ protocol (**Chen** *et al.*, 2013; **Scheres**, 2012)). Thus, with the exception of the least common subclass (v), the class of complete ribosomes with a complete set of r-proteins displays tRNA occupancies and conformational variability compatible with active translational activity.

For the population of incomplete 80S ribosomes identified in the first classification step, further subclassification revealed five different subpopulations, distinguished by the presence versus absence of one or two r-proteins as well as conformational changes (Fig. S1 **and S2 and Table S3**): (a) 80S lacking density for RPL10 (uL16) (**Fig. 2A**); (b-c) 80S lacking densities for both RPL10 (uL16) and RPS1A/B (eS1) (Fig. 2B **and C**); (d-e) 80S lacking densities for both RPL10 (uL16) and RPS1A/B (eS1), as well as densities for small fragments of rRNA (Fig. 2D **and E**). The resolutions of these density maps range from 6.2Å to 9.8Å.

**Fig. 2.**
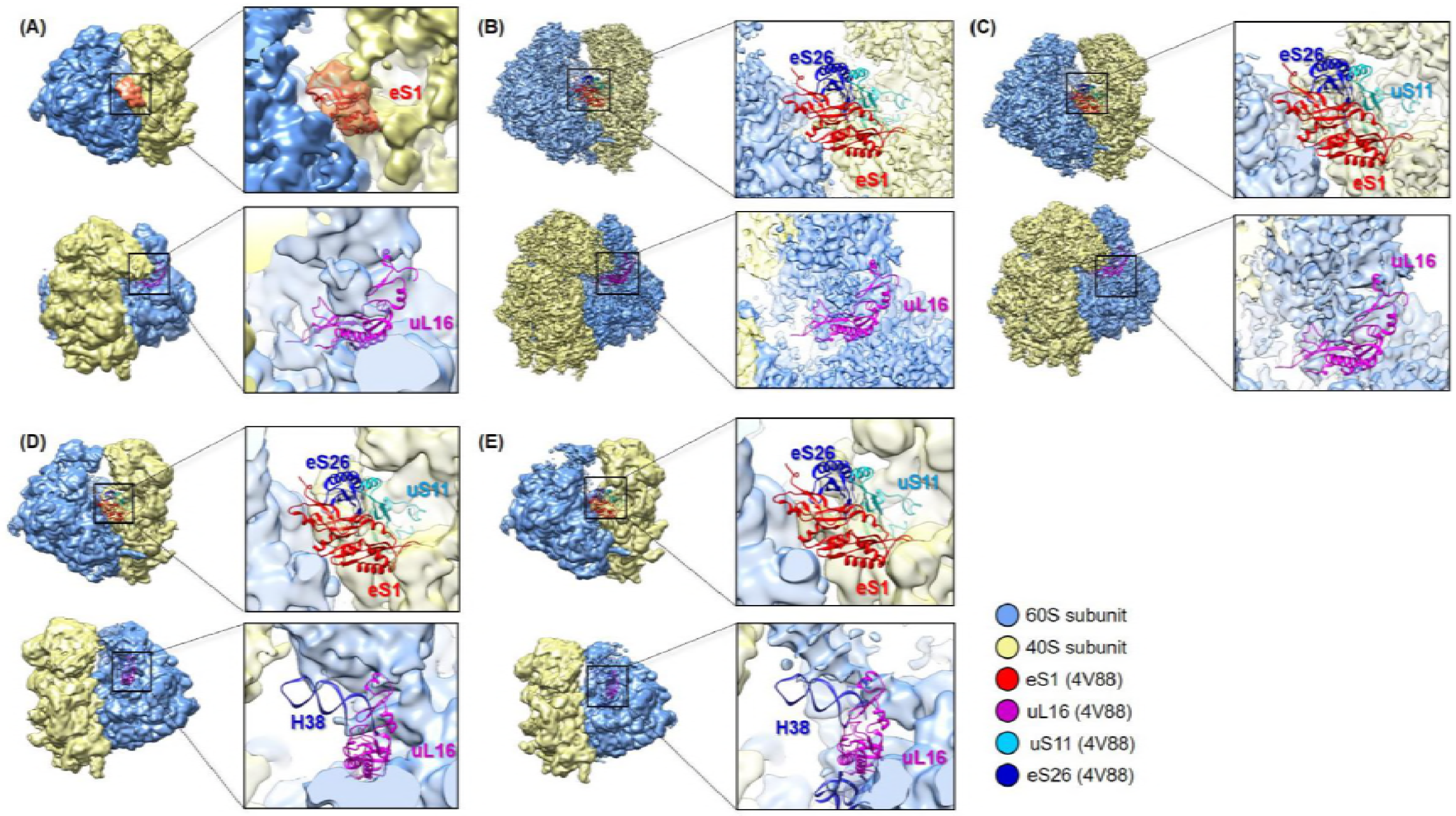
Cryo-EM reconstructions of *S. cerevisiae* 80S ribosomes after the switch from glucose to glycerol. Yellow, 40S subunit; blue, 60S subunit. In all panels. **A-E**, two different views of the ribosome are shown, related by a ~90-degree rotation around a vertical axis in the plane of the figure. On the right hand side in each panel, a magnified slab of a region marked in the left view is shown, with densities shown in transparency. **(A)** Density map reconstructed from the sub-population of incomplete 80S ribosomes that lack only the density for RPL10 (uL16) protein but possess protein RPS1A/B (eS1) (orange). **(B)** Rotated incomplete 80S ribosomes that lack the density for both RPS1A/B (eS1) and RPL10 (uL16) r-proteins, as well as the interacting partners of RPS1A/B (eS1),RPS26A/B (eS26), and RPS14A/B (uS11). **(C)** Non-rotated incomplete 80S ribosomes lacking the densities for both RPS1A/B (eS1) and RPL10 (uL16) r-proteins. **(D, E)** Incomplete 80S ribosomes lacking densities for both the RPS1A/B (eS1) and RPL10 (uL16) proteins, as well as the H38 of the 28S rRNA. The interacting partners of RPS1A/B (eS1) (RPS26A/B (eS26) and RPS14A/B (uS11)) are also missing entirely. The x-ray structures of *S. cerevisiae* (PDB 4V88) 40S and 60S subunits were rigid-body fitted separately into cryo-EM densities using *UCSF Chimera* (**Pettersen** *et al.*, 2004).

### Locations and interacting partners of proteins uL16 and eS1

The most notable finding in our cryo-EM study is that upon carbon source switch from glucose to glycerol, the fraction of the 80S ribosome particles missing the r-proteins RPS1A/B (eS1) (from 40S subunits) and RPL10 (uL16) (from 60S subunits) increases rapidly. The small-subunit r-protein RPS1A/B (eS1) is located in the vicinity of the mRNA exit tunnel and has direct contacts with helix 26 (H26) of the 18S rRNA and r-proteins RPS14A/B (uS11) and RPS26A/B (eS26). In the class of incomplete 80S ribosomes lacking RPS1A/B (eS1), we also found that RPS14A/B (uS11) and RPS26A/B (eS26) either displayed scattered densities or were completely absent from the ribosome, suggesting weak binding affinities of these proteins to 40S subunits lacking RPS1A/B (eS1) (**Fig. 2**).

The large-subunit protein RPL10 (uL16) is located in the inter-subunit corridor through which tRNAs move during the peptide elongation cycle. It has contacts with several key, functionally important sites via its C-terminal region (P-loop), including the elongation factor binding site, the peptidyl-transferase center (PTC), and H38 of the 28S rRNA (also known as the A-site finger). Protein RPL10 (uL16) also has interactions with the 5S rRNA via its C-terminus and with H69 of the 28S rRNA via its conserved internal loop (**Sulima** *et al.*, 2014). In the population of incomplete 80S ribosomes, we found that H38 and its neighboring helices display only scattered densities (Fig. 2D **and 2E and Table S3**), indicating variable positions. Such positional variability is not commonly observed in mature 80S ribosome structures but has been found in another, independent study of 60S subunit biogenesis (**Malyutin** *et al.*, 2017).

### The binding affinity of eS1 and uL16 to 80S ribosomes changes upon switch of carbon source

To quantitatively investigate the binding behavior of RPS1A/B (eS1) and RPL10 (uL16) to ribosomes, we monitored the number of ribosomal particles that were assigned to the populations of ‘complete 80S’ and ‘incomplete 80S’ at different time points (T = 0, 30, 120, 240, 450, and 1440 min after changing growth in glucose to glycerol) using our pooled classification strategy. In order to test experimental reproducibility, we performed the same pooling and tracking strategy in two independent cryo-EM experiments. In both studies, the quantity of particles assigned to the ‘incomplete 80S’ population increased in the first 30 min after shifting from glucose to glycerol, as the binding ratio of RPL10 (uL16) and RPS1A/B (eS1) to 80S ribosomes decreased from ~83% to ~66%, and then remained relatively stable: ~64% at T=120 min and ~62% at T=240 min (**Fig. 3**). After incubation in glycerol for 450 min, the binding of RPL10 (uL16) and RPS1A/B (eS1) to 80S ribosomes recovered from 62% to ~71% at T=450 min and stabilized at ~73% at T=1440 min (24 h) (**Fig. 3**).

**Fig. 3.**
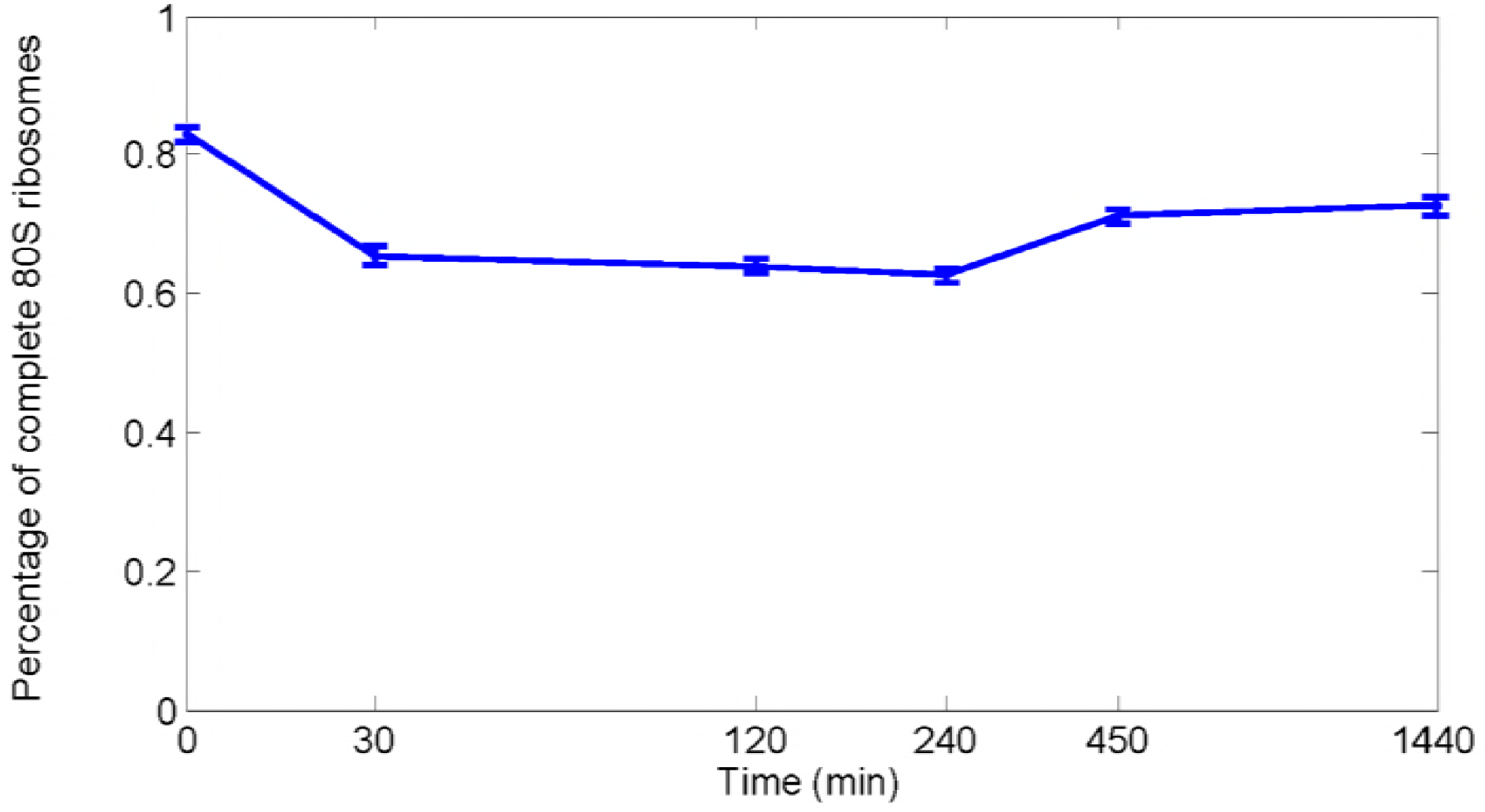
Structural inventory as a function of time. The x-axis represents the time (min, log scale) that cells were incubated in glycerol medium. The y-axis represents the percentage of complete 80S ribosomes. Generally, the percentage of complete 80S ribosomes in each sample set decreased upon onset of the glucose-to-glycerol switch. This change reflects the general decrease in binding of RPS1A/B (eS1) and RPL10 (uL16) to 80S ribosomes.

### Quantitative proteomic analysis of r-proteins in purified ribosomes

To identify and measure the changes in the protein composition of the same ribosome populations purified from yeast growing in glucose or glycerol, we analyzed the ribosomal protein composition using iTRAQ-labeling and quantitative tandem mass spectrometry (**Link** *et al.*, 1999; **Ross** *et al.*, 2004). Whole-cell extract and purified ribosome sample from yeast cells grown in glucose (T=0) and the 24h glycerol (T=24h) time points were used. After proteolytic digestion of the proteins with trypsin, the N-termini and amino acid side chain amines of the tryptic fragments were covalently labeled with isotoptic 4-plex mass tags (**Ross** *et al.*, 2004; **Unwin** *et al*., 2010). After replicate ribosome samples from the glucose (T=0) and the 24h glycerol (T=24h) time points were labeled with unique iTRAQ tags, we pooled the iTRAQ-labeled ribosome samples and fractionated the peptide mixture using multidimensional microcapillary HPLC (**Link** *et al.*, 1999). The eluting peptides were analyzed using nanoESI-tandem mass spectrometry (MS/MS)(**Samir** *et al.*, 2015). The peptide fragmentation data were computationally compared to predicted values from a yeast protein database to identify the labeled peptides and hence the corresponding proteins (**Eng** *et al.*, 1994). The fragmentation of the iTRAQ tag generated low molecular mass reporter ions that were used to quantify the identified r-protein peptides and proteins (**Ross** *et al.*, 2004; **Unwin** *et al*., 2010).

From three independent biological replicates, we identified 135 of the 139 r-proteins from the purified ribosome complexes using mass spectrometry-based proteomic (**Table S4**). Of the 79 r-yeast proteins in the 80S ribosomal complexes, we successfully identified 20 of the 22 r-proteins from single-copy genes and 55 of the 57 r-protein pairs from paralogous genes. The four proteins not identified were from r-protein genes encoding small proteins < 60 amino acids in size. The mass spectrometry analysis identified 21 of the 22 100% identical paralogous r-proteins (**Table S4**). For 16 of the 35 paralogous gene pairs that are not 100% identical, we successfully identified both unique peptides. Of the remaining 19 non-identical paralogous gene pairs, our mass spectrometry analysis was able to uniquely identify 1 of the 2 paralogous r-proteins in 8 cases (**Table S4**).

For each of the three replicates, we next calculated the relative fold-change in abundance for 131 r-proteins in glycerol (T=24) compared to glucose (T=0) using the log_2_ ratio of the iTRAQ reporter ion signal intensities (**Table S5**). Because ribosomes were previously purified and analyzed from identical whole-cell extracts (**Samir** *et al.*, 2015), we were able to directly compare the fold changes of 126 r-proteins quantitated in both the whole-cell extracts and purified ribosomes. A correlation analysis between the fold change of r-proteins in the solubilized whole cell extracts and the purified ribosome extracts shows poor correlation (Fig. 4A **and B**). However, there was strong correlation in the fold-ch nges of r-proteins between the experimental replicates (Fig. 5A **and B**). The lack of correlation between the fold changes in cell extracts and purified ribosomes suggest that a fraction of r-proteins in the cell are not incorporated into ribosomal complexes. We reasoned that either these r-proteins have not yet been successfully incorporated into ribosomal complexes during ribosome biogenesis or ribosomes were disassembling.

**Fig. 4.**
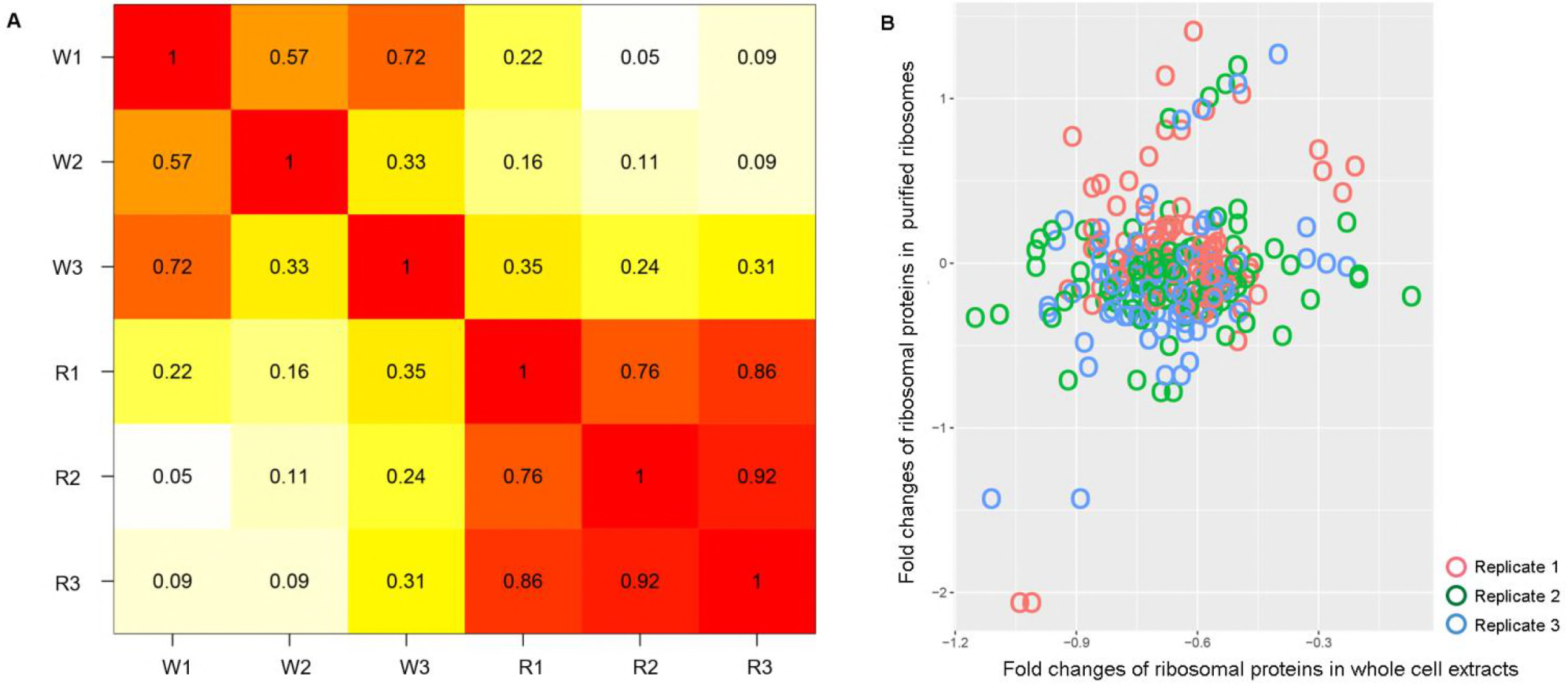
Correlation analysis between the fold changes of r-proteins in whole cell extracts and purified ribosomes from cells grown at standard conditions (glucose, 30°C) and glycerol (glycerol, 30°C).(See Tables S2 and S3) The summary of three replicates is shown. **(A)** Cross-correlation matrix. Numbers represent Pearson’s R. W# represent whole extract replicates. R# represent purified ribosome replicates. **(B)** Scatter plot showing the relationship between fold-changes in whole cell extracts (x-axis) and purified ribosomes (y-axis).

**Fig. 5.**
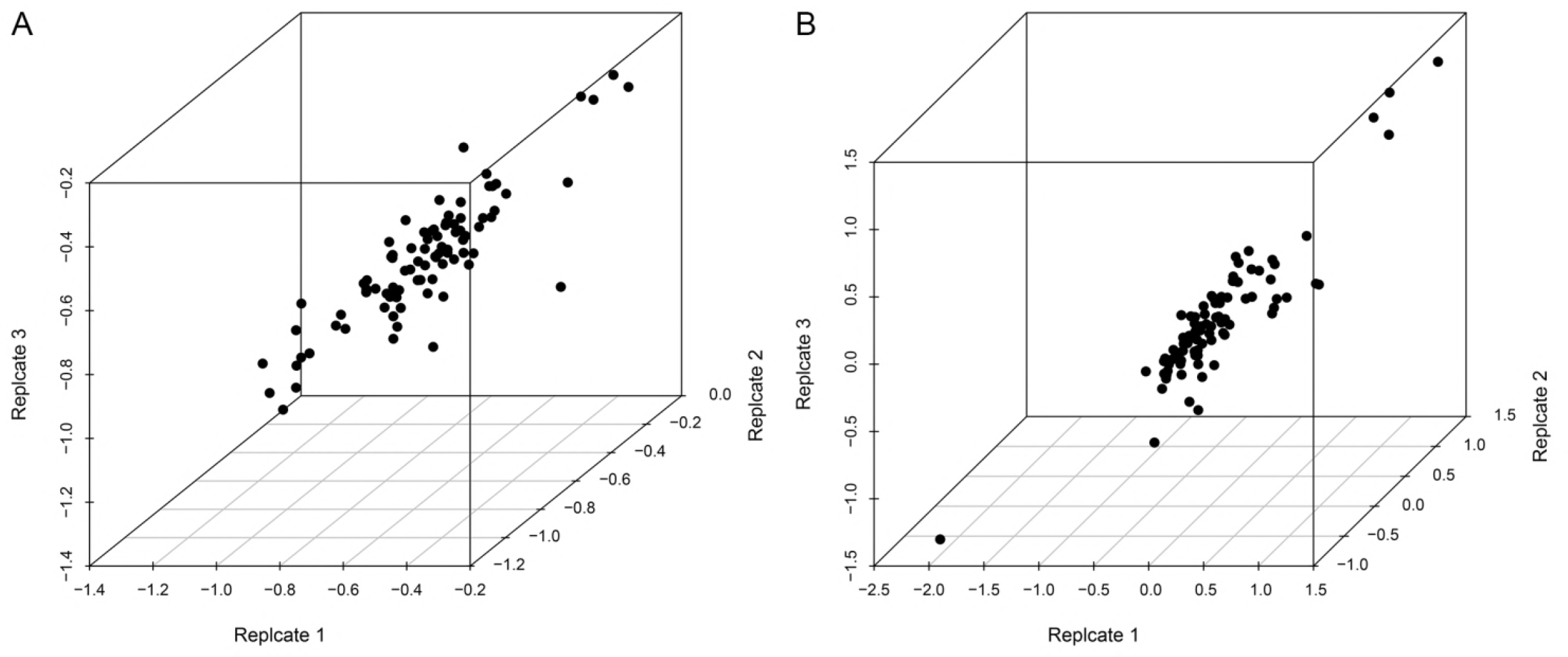
Scatterplots showing reproducibility among replicates of whole cell extracts and purified ribosomes. **(A)** Whole-cell extract replicates. **(B)** Purified ribosome replicates. Plots were generated in *RStudio*.

To identify differentially abundant r-proteins in purified yeast ribosomes grown in glucose and glycerol, we used a *t-*test of independence with an alpha level of 0.05. Since there are only minimal sequence differences between the r-protein paralogs to reliably quantify the paralog-specific changes, we reanalyzed the mass spectrometry data using only the unique peptides for quantitation. We quantitated 45 r-proteins with at least one unique peptide (**Fig. 6A**), and identified 11 r-proteins that were differentially present in the purified ribosome complexes from yeast cells grown in glucose or glycerol (**Fig. 6B**). The list of significantly changing r-proteins in the purified ribosomes contained the paralog pair RPL8A (eL8A) and RPL8B (eL8B) (97% protein sequence identity) whose fold changes were opposite to each other. To validate the iTRAQ data, the relative difference in abundance of RPL8A (eL8A) and RPL8B (eL8B) proteins in the purified ribosomes was measured using multiple reaction-monitoring mass spectrometry (**Fig. 6C**). The data suggest yeast cells have a differential requirement for RPL8A and RPL8B under the two growth conditions.

**Fig. 6.**
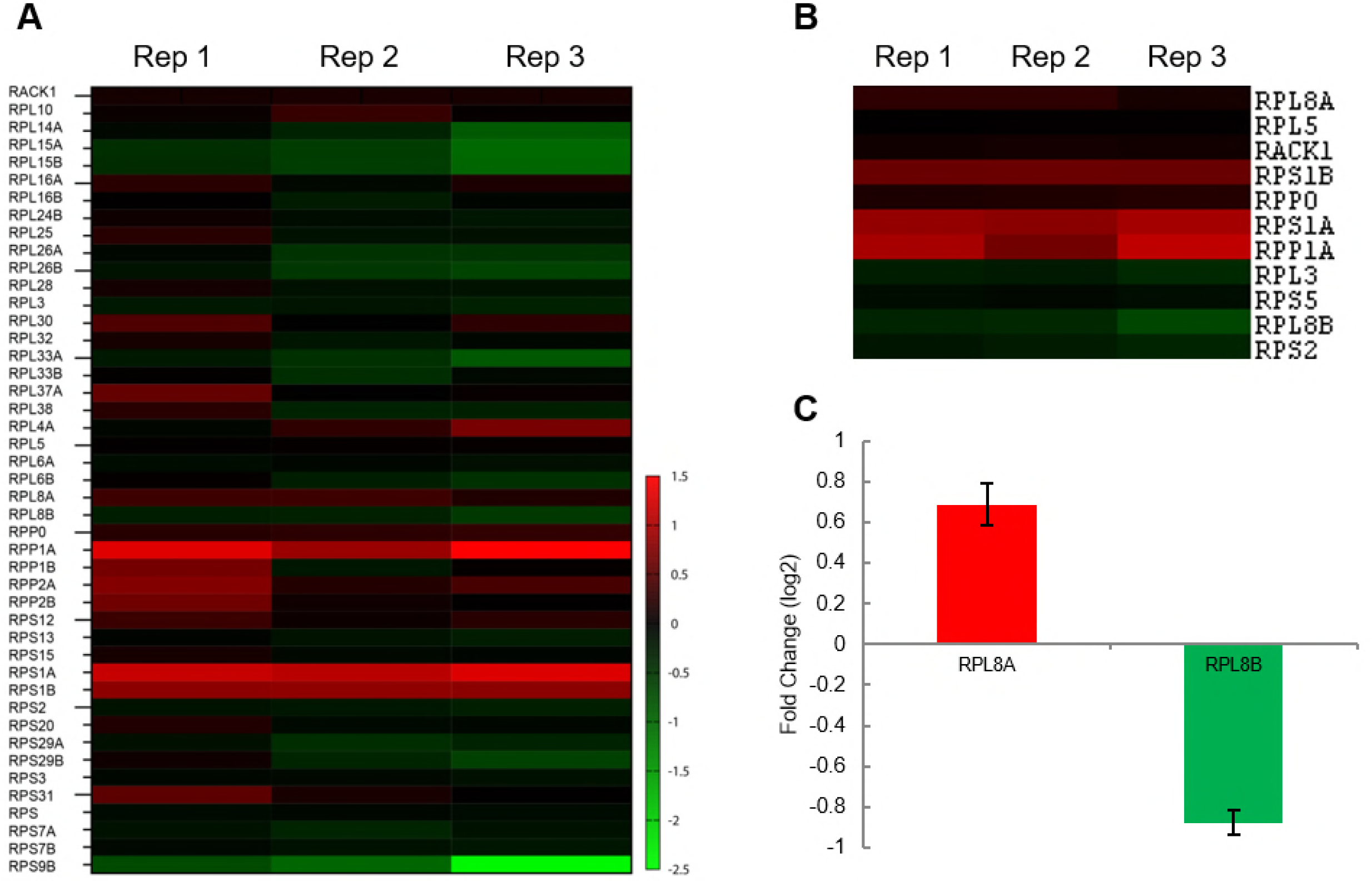
Quantitation of r-proteins in ribosomes purified from yeast cells grown in glucose and glycerol. **(A)** Forty-five ribosomal proteins identified and quantitated using at least one unique peptide in iTRAQ-mass spectrometry experiments. The log_2_ transformed fold change in the r-protein’s relative abundance 24 h after changing growth in glucose to glycerol was used to generate the heatmap. **(B)** Eleven statistically significant differentially abundant r-proteins in the purified ribosomes from cells grown in glucose and glycerol identified using a t-test’s *p*-value less than 0.05. **(C)** The protein abundance of RPL8A and RPL8B validated using multiple reaction monitoring in three independently purified ribosome replicates. Color Legend: red increased; green decreased abundance of the yeast r-protein in cells growing in glycerol compared to glucose media.

The cryo-EM data showed ~83% to ~66% of the 80S ribosomes from the purified ribosomal complex missing r-proteins RPS1A/B(eS1) and RPL10(uL16). MS-proteomics successfully identified both r-proteins (**Table S5**). However, the quantitative proteomic analysis showed both proteins increasing their relative abundance in the purified ribosomes 24 h after changing from glucose to glycerol media, although the change in RPL10 abundance was not statistically significant **(Fig 6A-B**). To explain the discrepancy, we suspect the sucrose gradient-purified 80S ribosomes contained both 80S ribosomes as well as sub-80S particles including 40S, 60S, and partially assembled cytoplasmic ribosomal complexes. Our cryo-EM computational image analysis was able to exclude ribosomal complexes smaller than 80S particle while our MS-proteomics analysis could not. We cannot exclude the possibility that Rps1A/B and RPL10 become post-translationally modified when changing growth from glucose to glycerol, thereby distorting our quantitative proteomics analysis. Although the 80S ribosomal samples used in the cryo-EM and proteomics experiments were the same, the two analytical methods and resulting computational analysis likely targeted different classes of ribosomal complexes, limiting our ability to directly compare the cryo-EM and quantitative proteomic data.

### Change of polysomal fractions in response to switch of carbon source

Since we found that with the shift to glycerol, the percentage of complete 80S ribosomes decreased and their r-protein composition changed, we speculated that global translational activities might be affected. We performed polysome profiling experiments using the same time course. These experiments indeed revealed a re-distribution of polysomes (actively translating) into the 80S peak (inactive) after 30 min following the shift from glucose to glycerol (**Fig. 7**). After 450 min, the polysomes partially recovered (**Fig. 7**). A similar re-distribution of polysomal fractions to the 80S fraction was also observed after cells were shifted from glucose to glycerol for 10 min (**Kuhn** *et al.*, 2001).

**Fig. 7.**
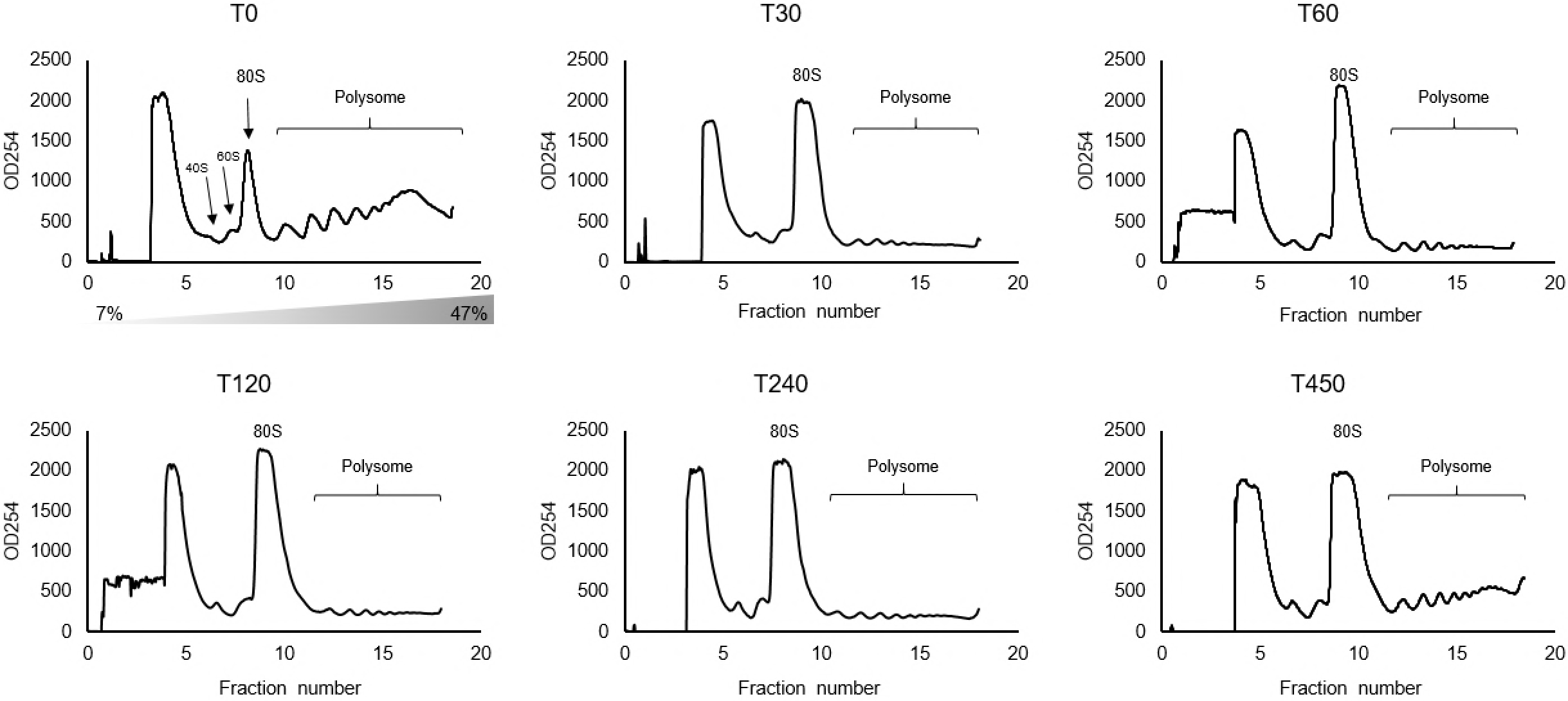
The switch of carbon source from glucose to glycerol inhibits translation. Polyribosome traces from the wild-type *S. cerevisiae* strain. Yeast was grown in complete medium containing glucose, then re-suspended at T=0 in medium containing glycerol for the indicated times (min). Polyribosomes were analyzed as described in ‘STAR Methods’. The peaks that contain the small 40S ribosomal subunit, the large 60S ribosomal subunit, and complete 80S ribosomes are indicated by arrows. The polysome peaks are bracketed.

The cryo-EM data showed ≈20-40% of the yeast 80S ribosomes rapidly lose RPL10 and RPS1A/B subunits within 30 min after the change from glucose to glycerol. Because ribosome biogenesis in eukaryotes is a considerably longer process involving multiple steps and interactions, it is highly unlikely that new ribosomes are being formed within 30 min. We reasoned the cell’s 80S ribosomes are losing r-protein subunits and adopting an inactive state when shifted from glucose to glycerol. Since the cryo-EM classification method employed is unable to differentiate between 80S ribosomes containing RPL8A (eL8A) and RPL8B (eL8B), we do not know how the subpopulation of ribosomes lacking RPL10 (uL16) and RPS1A/B (eS1) is related to the one exhibiting the change from one r-protein paralog to the other. RPL8A and RPL8B. Future experiments are needed to confirm whether existing 80S ribosomes are exchanging RPL8B with cytoplasmic RPL8A or new ribosomal synthesis is required to change these r-protein paralogues in the cell’s population of ribosomal subunits.

### Different functional Roles of *RPL8A* and *RPL8B*

Using growth assays, we tested the paralog-specific roles of *RPL8A* and *RPL8B* using yeast genetics. When we compared yeast growth rates in glucose versus glycerol, we observed a functional difference in the activity or specificity of the yeast ribosome depending on the presence of eL8A and eL8B **(Fig. 8)**. Under optimal growth conditions in rich media with glucose as the carbon source, the function of either *rpl8a*Δ or *rpl8b*Δ null allele can be compensated for by the paralogous gene (**Fig. 8A**). However, when the cells encounter non-optimal growth in glycerol, the differential requirement for each paralog becomes apparent (**Fig. 8B-D**). This result is consistent with the original ribosome filter hypothesis in which the ribosome filter was proposed to fine-tune gene expression (**Mauro** and Edelman, 2002; **Mauro** and Edelman, 2007).

**Fig. 8.**
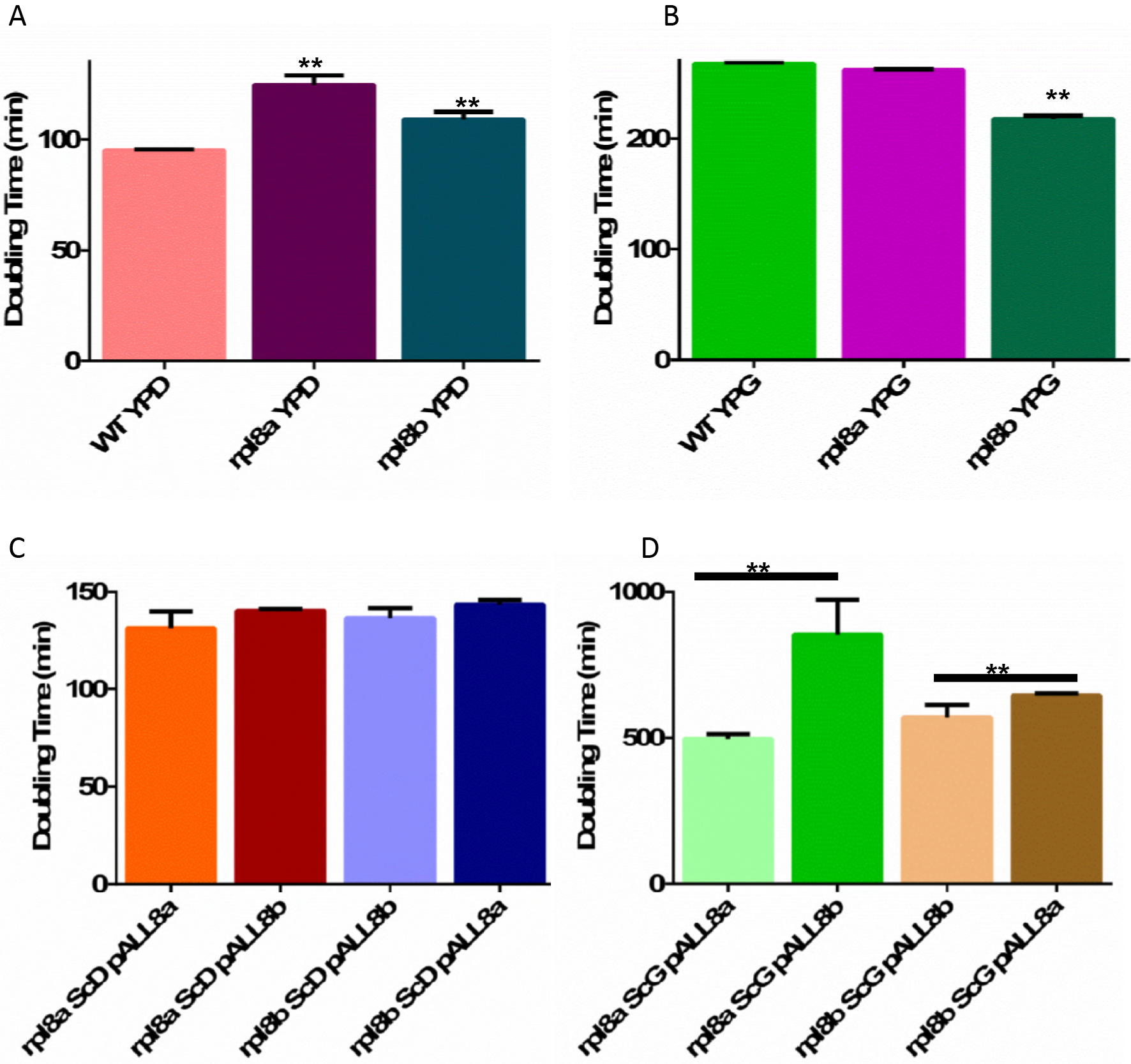
Doubling times comparisons. A & B) Stars denote the statistical significance of comparison with the WT strain (*p*-values < 0.05). A) Doubling times of WT, *rpl8a*, and *rpl8b* with glucose as carbon source. B) Doubling times of WT, *rpl8a*, and *rpl8b* with glycerol as carbon source. C) Doubling times of *rpl8a* with Rpl8a on plasmid, *rpl8a* with Rpl8b on plasmid, *rpl8b* with Rpl8b on plasmid, and *rpl8b* with Rpl8a on plasmid with glucose as carbon source. D) Doubling times of *rpl8a* with Rpl8a on plasmid, *rpl8a* with Rpl8b on plasmid, *rpl8b* with Rpl8b on plasmid, and *rpl8b* with Rpl8a on plasmid with glycerol as carbon source. Stars denote the statistical significance of comparison denoted by the lines above the bar graph (*p*-values < 0.05).

## Discussion

In this study we have employed both cryo-EM experiments and quantitative mass spectrometry along with polysome profiling and genetic analysis to follow changes in the yeast ribosome composition as a function of time after a shift from growth in glucose to glycerol media. Using a novel quantitative approach combining pooled classification with 3D visualization, our cryo-EM experiments identified and tracked subpopulations of ribosomes lacking individual r-proteins. The cryo-EM experiments enabled us to identify changes in 80S structure, protein composition, and bound tRNAs after switching from growth in glucose to glycerol. Using classification tools, we could pre-process the cryo-EM images to select only 80S particles for in-depth analysis and exclude smaller ribosomal particles. The quantitative proteomic experiments enabled us to identify and quantify changing r-proteins, including paralogs, incorporated into yeast ribosome populations that would be impossible to identify with the current cryo-EM methodology. Isotopic labeling enabled us to perform precise quantitative measurements of individual r-proteins in the purified ribosomal complexes. Unlike cryo-EM’s 80S/60S/40S classification capability, the quantitative proteomic experiments analyzed mixtures of purified ribosomal complexes isolated from sucrose gradients. The r-proteins’ low molecular weight and high abundance of lysine and arginine residues limited the number of tryptic peptides that could be surveyed by our mass spectrometry approach. Therefore, the number of r-proteins that could be confidently identified and quantified by the quantitative mass spectrometry was limited. Targeted mass spectrometry approaches would be one solution to this limitation. Overall, combining the two approaches of cryo-EM and mass spectroscopy allowed us to identify changing 80S ribosome compositions, structure, and functional status that would escape detection if only a single approach was employed.

Our identification of ribosomes with sub-stoichiometric compositions of r-proteins presents intriguing possibilities. The data analysis resulting in reconstructions of selected classes uncovered a sizeable fraction of translation-incompetent yeast ribosomes lacking both RPS1A/B (eS1) and RPL10 (uL16) in cells growing in glucose. Remarkably, this fraction almost doubled immediately (within 30 min) after the shift from glucose to glycerol. Apparently, an increasing number of the r-proteins RPL10 (uL16) and/or RPS1A/B (eS1) dissociate from existing complete ribosomes when yeast cells are grown in glycerol. Depletion of the two r-proteins from the ribosome could be functioning as a posttranscriptional regulatory mechanism to slow the overall rate of protein synthesis as both genes are essential in yeast (**Arevalo** and Warner, 1990). It is important to note that the presence of both proteins in the ribosome is essential for translation. RPS1A/B (eS1) forms part of the protein-protein and r-protein-rRNA contacts between two adjacent 40S subunits in polysomes (**Myasnikov** *et al.*, 2014), and RPL10 (uL16) plays a central role in the interaction of the ribosome with GTP-bound translation factors. The rapid loss of RPL10 (uL16) and/or RPS1A/B (eS1) from the 80S ribosome within 30 min after the change from glucose to glycerol suggests that the environmental change induces the loss of these two r-proteins from the existing pool of complete ribosomes, rather than the production of specialized ribosomes lacking these two proteins during the process of ribosomal biogenesis (**Gasch** *et al.*, 2000). While environmental changes, such as the switch from glucose to glycerol, induce rapid changes in yeast transcriptional profiles (**Gasch** *et al.*, 2000), ribosome biogenesis is a relatively slow process requiring multiple steps and factors (**Woolford** and Jonathan, 1991). Surprisingly, the proportion of complete, translation-competent ribosomes does not recover to the prior state. Since yeast cells grow slowly with glycerol as a carbon source compared to glucose, the ribosomes lacking both of the r-proteins may be part of a non-translating reserve ribosome pool (**Samir** *et al.*, 2015).

## Author Contributions

A.J.L. conceived the hypothesis. M.S., P.S., B.S., J.F., and A.J.L. designed the experiments. M.S. and B.S. performed cryo-EM experiments. M.S., B.S., and W. L. analyzed the cryo-EM results. P.S. and C.M.B. performed the iTRAQ, liquid chromatograph mass spectrometry and polysome profiling experiments. P.S. and Rahul analyzed the MS results. M.S., P.S., J.F. and A.J.L. wrote the manuscript. All authors approved the final manuscript.

## Acknowledgements

This study was supported by HHMI and NIH R01 GM29169 (to J.F.) and NIH RO1 grant GM64779 (to AL). CB was supported by NIH training grant T32 AI007611.

---

**Figure S1.**
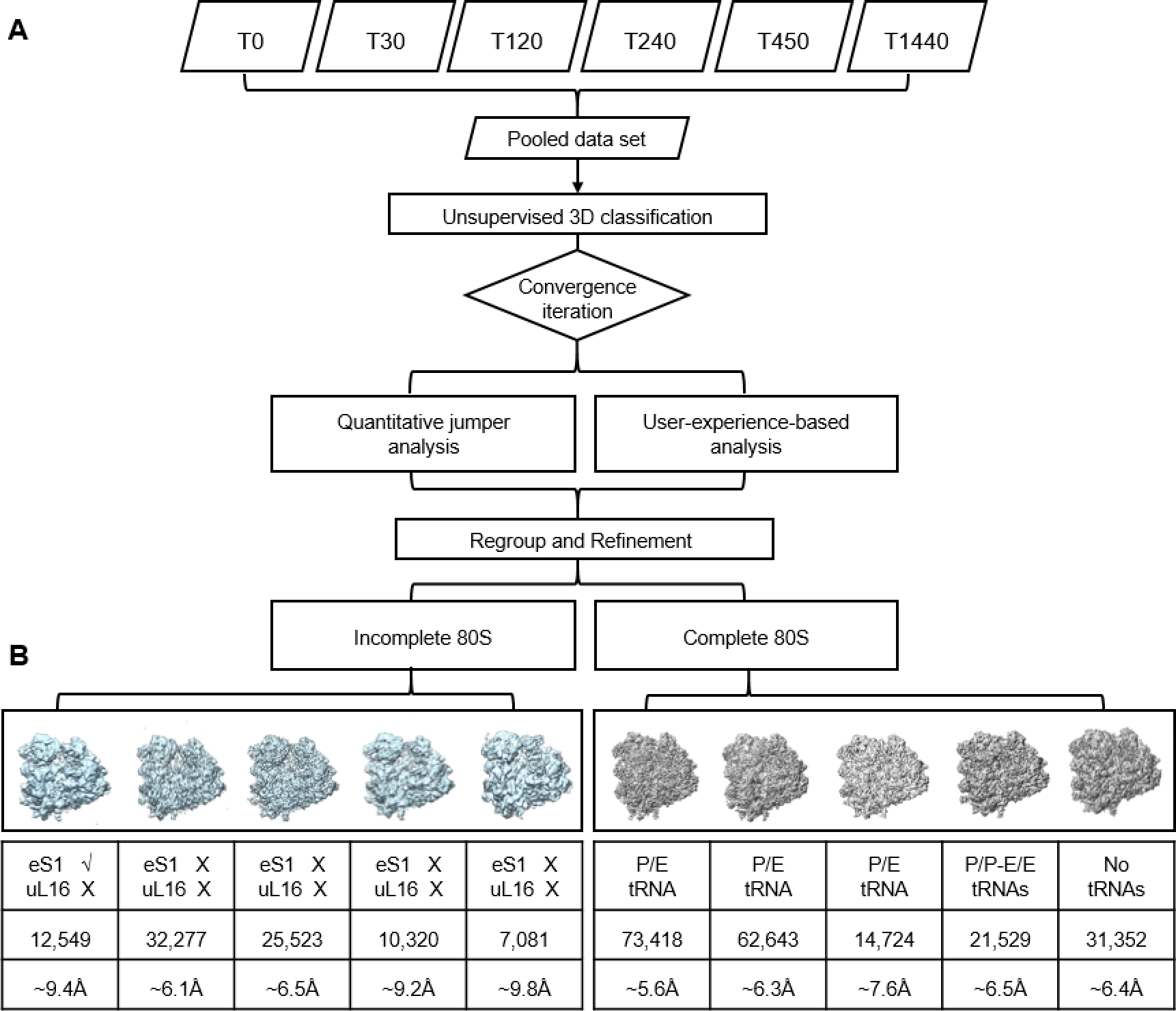
Full framework of ‘pooled’ classification strategy and results (Related to Figure 5) (A) Schematic diagram of our ‘pooled classification’ data processing procedure. Samples purified from six time points were pooled together and classified using the same procedure. Further analysis was based mainly on quantitative jumper analysis (20) with minimum user interference. (B) Summary of the full-scale cryo-EM studies. “√”, present; “X”, absent. Cryo-EM reconstructions of incomplete 80S ribosomes are presented in front view and colored in light blue (B, left), while complete 80S ribosomes are colored in grey (B, right). Resolutions reported are based throughout on the ‘gold standard’ protocol along with the FSC = 0.143 criterion, and involved soft masking and high-resolution noise substitution (Chen et al., 2013).

**Figure S2.**
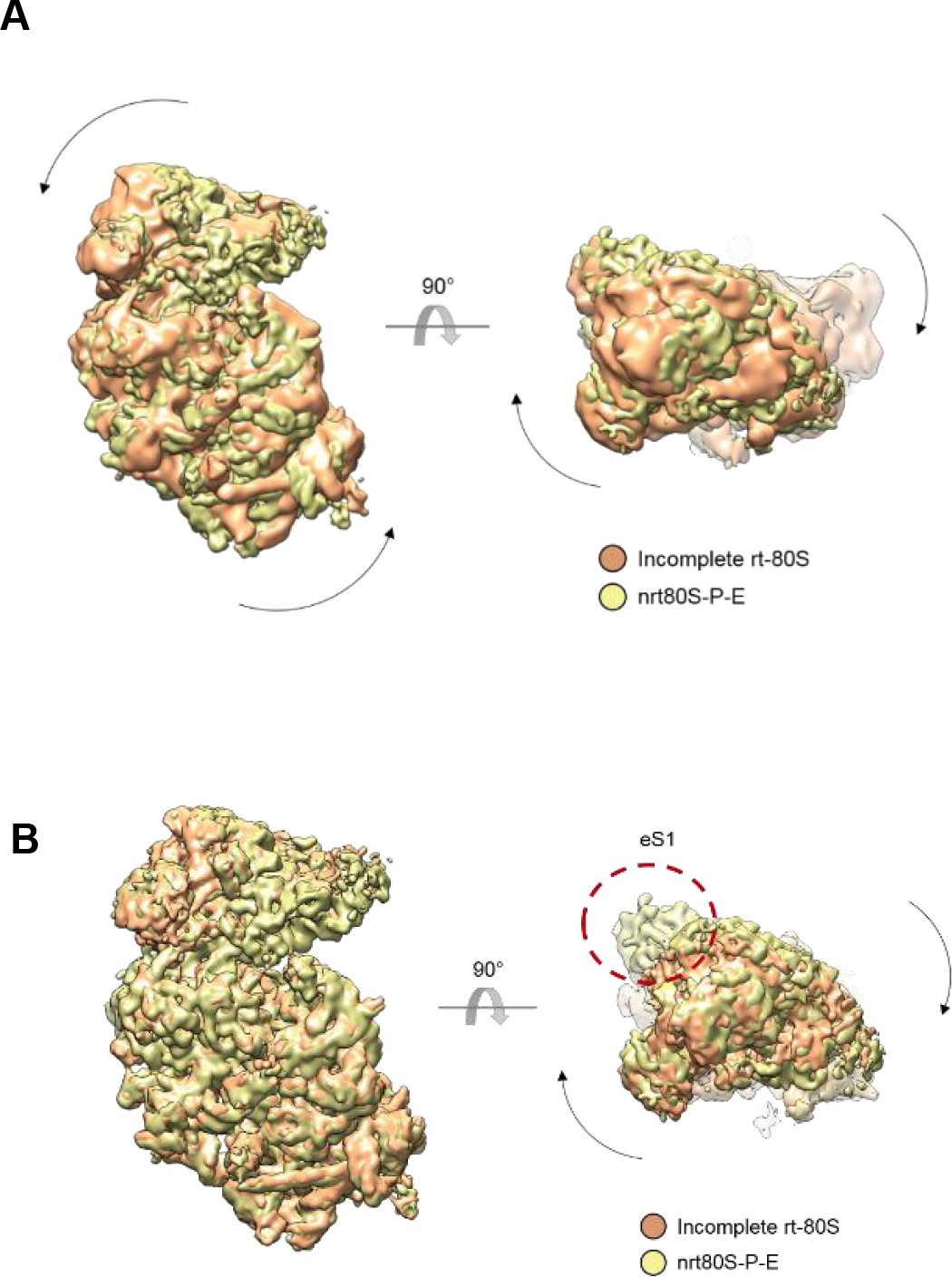
Global conformational changes of incomplete 80S ribosomes (Related to Fig 2) **(A)** Inter-subunit rotation and 40S subunit head-swiveling movements in the incomplete 80S (class 1). Comparison of the 40S subunit positions in nrt80S-P-E state (yellow) and the incomplete 80S (class 1) (orange). **(B)** Head-swiveling movements in the incomplete 80S (class 3). The position of eS1 on nrt80S-P-E state is highlighted by a red circle. Comparisons were obtained by structural alignment on the 60S subunits of the 80S ribosomes using UCSF Chimera (Pettersen et al., 2004).

**Supplemental Table S5:**
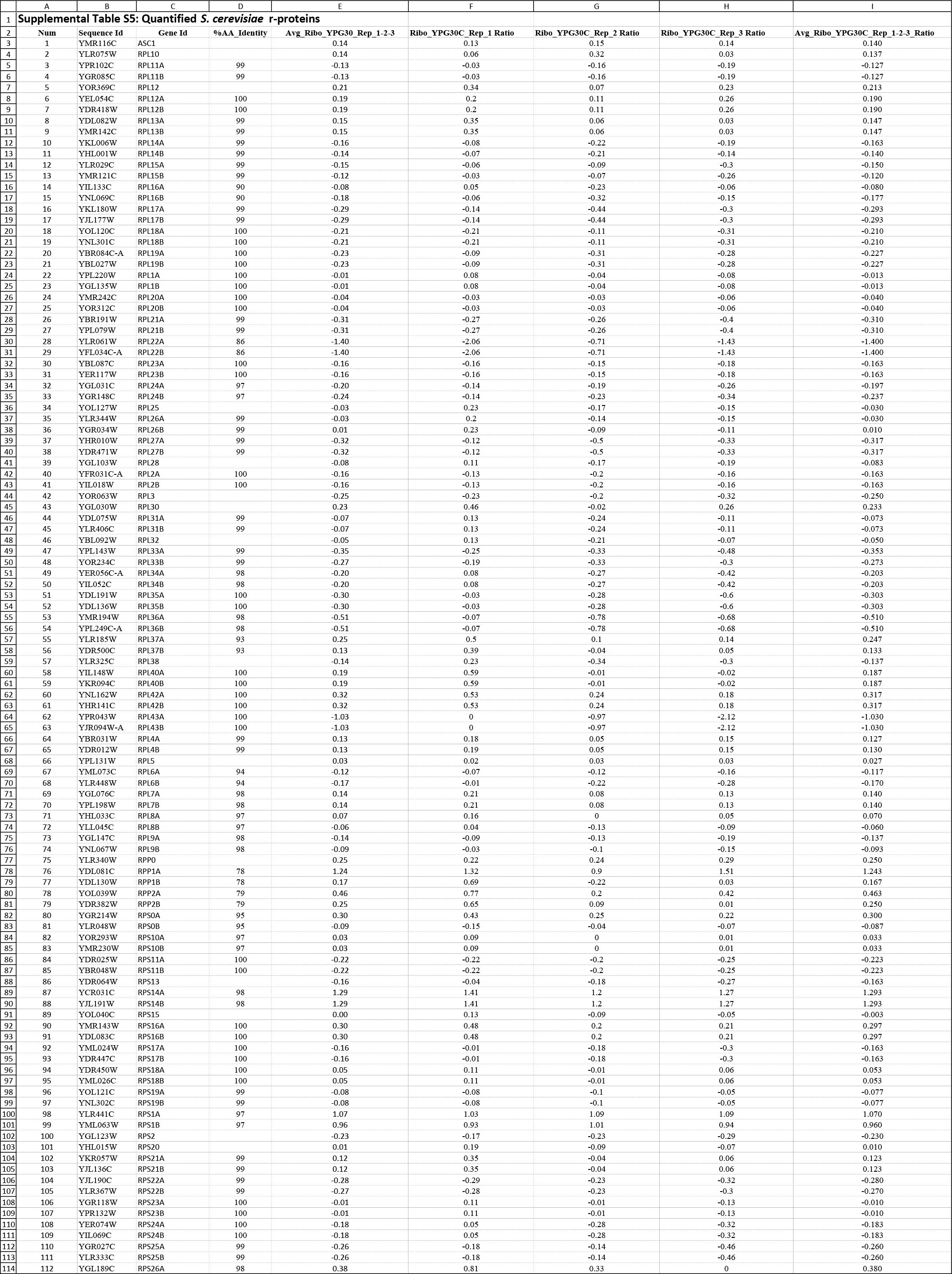
Quantified *S. cerevisiae* r-proteins

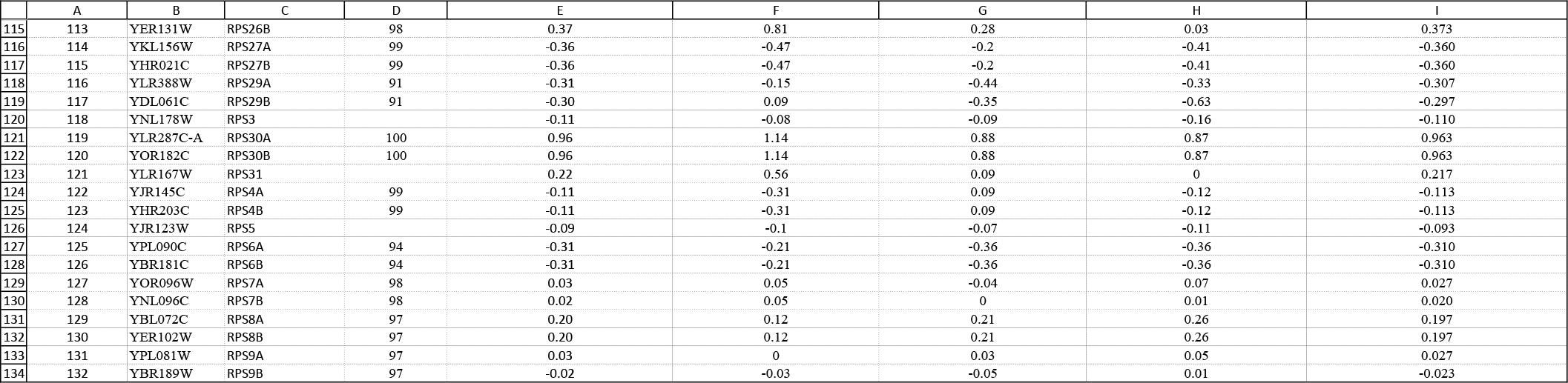

## Methods

### Data and Software Availability

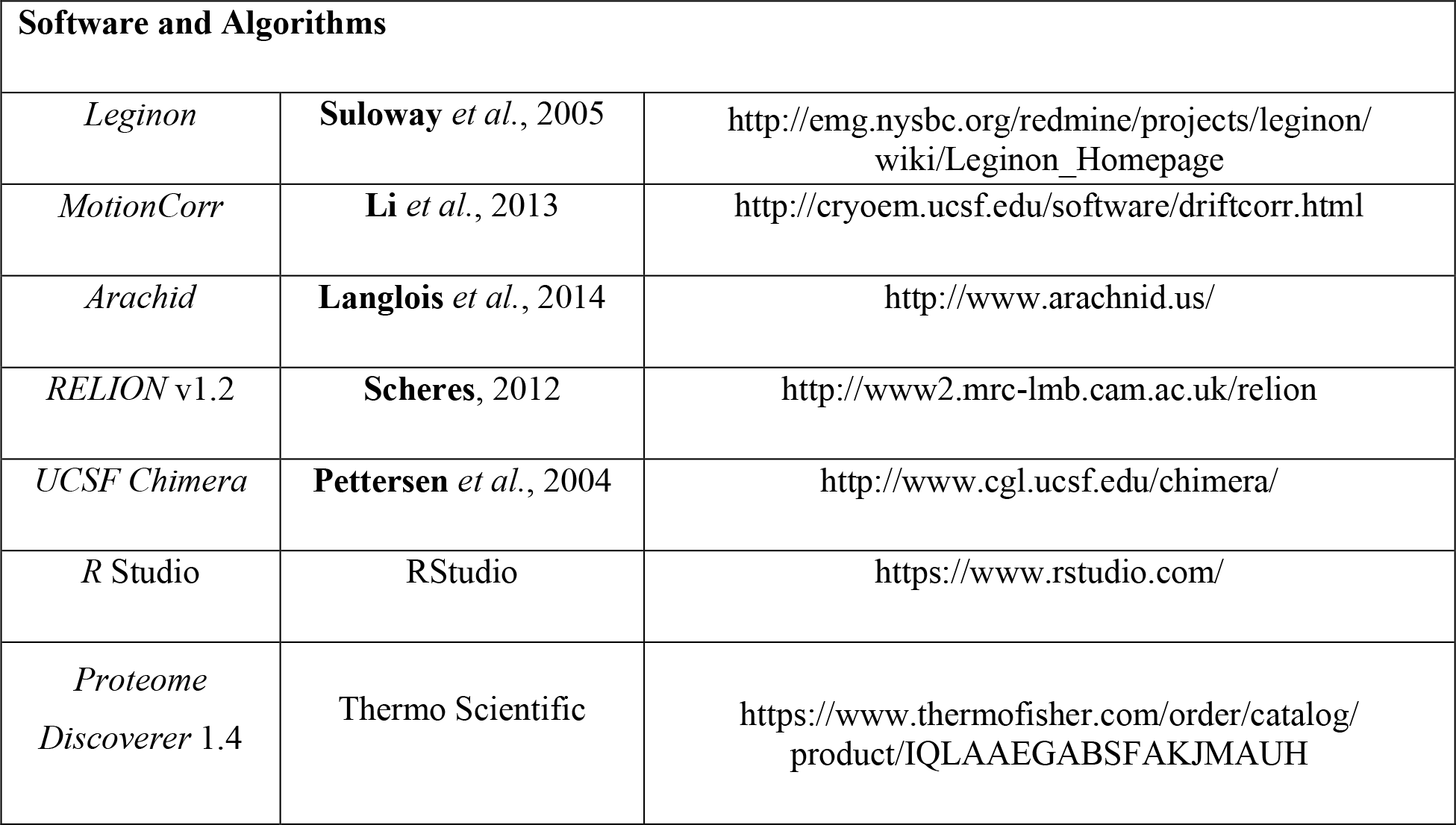

### Contact for Reagent and Resource Sharing

Requests for reagents may be directed to the lead contacts Andrew J. Link and Joachim Frank

### Experimental Model and Subject Details

#### Strains and Media

All experiments used the diploid *S. cerevisiae* strain BY4743, which has been previously described (**Brachmann** *et al.*, 1998). Cells were grown using standard techniques (**Amberg**, 2005). The genomic region of RPL8a and RPL8b, including ~1kb upstream and ~200b downstream, were cloned into the pFA6a-His3MX6 plasmid with Gibson cloning (New England Biolabs) to generate plasmids pALL8a and pALL8b, respectively (**Gibson** *et al.*, 2009). Strains were transformed using the standard yeast transformation protocol (**Amberg** *et al.*, 2005).

For calculating doubling times in YPD and YPG media, 5 mL starter cultures from a single colony of BY4743 (WT), *rpl8a*Δ(rpl8a), and *rpl8b*Δ (rpl8b) null strains were grown for 24 h as previously described (**Samir** *et al.*, 2015). For calculating doubling times of *rpl8a*Δ and *rpl8b*Δ strains complemented with plasmids pALL8a or pALL8b, yeast was grown in SC-His glucose (ScD) or SC-His glycerol (ScG) as previously described (**Samir** *et al.*, 2015; **Baker Brachmann** *et al.*, 1998; **Winzeler** *et al.*, 1999).

For plasmid rescue experiments, yeast strains were grown in SC–His supplemented with G418. One hundred microliters of the different media were added to wells in a 96-well plate as shown in Table S4. Cells were grown at 30°C in a Synergy Biotek plate reader for 720 min with constant shaking. OD 660 readings were taken every 3 min. A text file containing the OD 660 readings was exported and parsed using a custom *Python* script. Doubling times were calculated using the *lm* function in *R*. A Student’s *t*-test of independence was used to calculate the statistical significances at alpha level of 0.05. All *p*-values were adjusted for multiple hypotheses testing using the Bonferroni correction (**Dunn**, 1959, 1961). The source code used in this project is found in supplementary data.

### Method Details

#### Ribosome isolation and purification

*S. cerevisiae* cells were grown in YPD medium (1% yeast extract, 2% peptone, 2 % glucose) and shifted to YPG (1% yeast extract, 2% peptone, 3% glycerol). At the following time points ribosomes were isolated and purified: 0 min, 30 min, 120 min, 240 min, 450 min and 1440 min. Cells were first centrifuged at 2000 rpm for 5 min at 4°C using a Sorvall HLR6/H600A/HBB6 rotor in a Sorvall RC-3B centrifuge and washed with ice cold deionized H_2_O. The pellets were resuspended in 1 mL ice cold wash buffer (10 mM Tris pH 8, 5 mM β-mercaptoethanol, 500 mM ammonium chloride, 100 mM magnesium acetate) and lysed at 4°C for 10 min using glass beads and a Bead Beater (BioSpec, Inc) as previously described (**Browne** *et al.*, 2013). The cell suspensions were clarified by centrifugation at 20,000g for 15 min at 4°C. The pellets were discarded and the supernatants were overlaid onto 5-20% discontinuous sucrose gradients prepared in wash buffer. The gradients were centrifuged at 28,000 rpm using a SW-41 swinging bucket rotor for 18 h at 4°C. The ribosome-enriched pellets were resuspended in 1 mL ice cold standard buffer (10 mM Tris pH 8, 5 mM β-mercaptoethanol, 50 mM ammonium chloride, 5 mM magnesium acetate) and centrifuged for 10 min at 10,000g at 4°C. The pellets were discarded, and the ribosome suspensions were stored at −80° C.

#### iTRAQ labeling, liquid chromatography mass spectrometry, and data analysis

iTRAQ labeling, liquid chromatography tandem mass spectrometry, and database searches for peptide spectrum matching were done as previously described (**Eng** *et al.*, 1994; **Link** *et al.*, 1999; **Samir** *et al.*, 2015). *Proteome Discoverer 1.4* (Thermo Scientific) output files were imported into *ProteoIQ* (Premier BioSoft) for protein assembly and reporter ion quantitation. Peptide and protein FDR was set to 5%. Reporter ion normalization was based upon their intensities for the single copy r-proteins. This was done for both purified ribosome experiments and the previously published whole cell extract experiment (**Samir** *et al.*, 2015). The purified ribosome experiment was analyzed in two ways. For comparison with the whole cell extract analysis, all peptides were used for quantitation and protein assembly. To identify differentially present r-proteins, only the peptides uniquely mapping to one r-protein were used for quantitation and protein assembly.

Proteotypic peptides were selected for targeted quantitation from a database of identified peptides in the iTRAQ experiments. Transitions for unscheduled scout experiments were selected based upon NIST and GPM spectral libraries. Fifty μg of the purified ribosomes was digested with sequencing grade modified trypsin (1:50; Promega Corporation) and desalted essentially as described (**Samir** *et al.*, 2015). Peptides were eluted using an elution buffer composition of 50% acetonitrile, 0.1% trifluoroacetic acid. Peptides were analyzed using a 90 min scheduled SRM analysis. Briefly, peptides were autosampled onto a 200 mm by 0.1 mm (Jupiter 3 micron, 300A), analytical column coupled directly to an TSQ-Vantage (ThermoFisher) using a nanoelectrospray source and resolved using an aqueous to organic gradient (1-45% Buffer B) at a 500 nl/min flow rate. Using a series of unscheduled scout runs to determine retention times and transitions to monitor, a scheduled instrument method encompassing a 10 min window around each retention time along with calculated collision energies was created using *Skyline* (**MacLean** *et al.*, 2010). Q1 peak width resolution was set to 0.7, collision gas pressure was 1 mTorr, and the EZmethod cycle time was 5 s.

The resulting RAW instrument files were imported into *Skyline* for peak-picking and quantitation (**MacLean** *et al.*, 2010). The peak areas of the transitions were exported, and further analysis was done in Microsoft Excel. The sum of the peak areas of all the transitions of a given peptide, the peptide peak area, was used as the quantitative measure of abundance for the peptide. The average of peptide peak areas of all the peptides from a given protein, the protein peak area, was used as the quantitative measure of abundance of the protein. The average protein peak areas of the single copy r-protein RPL5p was used as a control. For differential analysis, in the first step, a ratio of peak area of the test protein to the peak area of the control was calculated across all samples. In the next step, a two sample *t*-test with alpha level 0.05 was performed with the ratios to test for statistical significance. Finally, the fold change was calculated by taking the ratio of the average of ratios. The calculated fold change was log2 transformed.

#### COMPzilla

*COMPzilla* uses CYC2008 2.0, a manually curated database of biomolecular complexes in yeast to identify complexes that are differentially present (**Pu** *et al.*, 2007). In the first step *COMPzilla* creates a dictionary with complex names as keys and the proteins that constitute the complex as values. Next, *COMPzilla* creates a dictionary in which complex names are still the keys, but values are mapped to fold changes of the proteins that constitute the complex. In the third step, *COMPzilla* creates a list all the fold changes in the population for statistical testing. Finally, *COMPzilla* compares the fold change distributions associated with protein complexes with population fold change distribution using a two sample *t*-test of independence and two-sample Kolomogorv-Smirnov test. *COMPzilla* exports the results of the two tests in separate tab delimited text files, in which the first column contains the complex names, the second column contains t-statistics or ks-statistics, and the third column contains the corresponding *p-* value. The *CompZilla.py* source code is available in the supplementary data.

#### Polysome profiling

Polysome profiling was modified from previous methods (**Browne** *et al.*, 2013). *S. cerevisiae* cells were grown in YPD to mid-log phase. Strains were then transferred to YPG and allowed to continue to grow. At 30 min, 120 min, 240 min and 450 min, cells were collected and lysed as previously described. Protein extracts were layered on top of 7% - 47% sucrose gradients and centrifuged.

#### Cryo-electron microscopy

Four microliters of purified ribosomes was applied to holey carbon grids (carbon-coated Quantifoil R2/4 grid, Quantifoil Micro Tools, GmbH, Groβlöbichau, Germany) containing an additional continuous thin layer of carbon, and glow-discharged using Gatan Solarus 950 (**Grassucci** *et al.*, 2007). Grids were blotted for 4 s at 4°C in 100% humidity and vitrified by plunging into liquid ethane cooled with liquid nitrogen, using the Mark IV Vitrobot (FEI, Hillsboro, Oregon) (**Dubochet** *et al.*, 1988).

For the initial studies, five data sets (T=0, 30, 120, 240, 450, and 1440 min) were collected on an FEI Tecnai F20 electron microscope (FEI, Hillsboro, Oregon) operating at 200 kV, equipped with a 4k x 4k CCD camera (Gatan, Pleasanton, CA). Images were recorded using the automated data collection system Leginon (Suloway *et al.*, 2005), and taken at the magnification of 5,000 x, corresponding to a calibrated pixel size of 2.245 Å.

For the full-scale study, data (T=0, 30, 120, 240, 450 and 1440 min) were collected on a TF30 Polara electron microscope (FEI, Hillsboro, Oregon) operating at 300 kV, set up with a K2 Summit direct electron detection camera (Gatan, Warrendale, PA). Images were recorded using the automated data collection system, Leginon (**Suloway** *et al.*, 2005), in counting mode, and taken at the nominal magnification of 23,000 x, corresponding to a calibrated pixel size of 1.66 Å. The dose rate was nominally set to 8 electron counts per physical pixel per second (**Liao** *et al.*, 2013), and the total exposure time was 8 seconds. Image stacks were collected in a defocus range of −1.5μm to −3.5μm and fractionated into 20 frames, each with an exposure time of 0.4 s.

#### Image processing

The six time-resolved sample sets were pooled and processed together. First, the dose-fractionated image stacks were corrected for beam-induced motion, using the method of Li *et al.* (**Li** *et al.*, 2013), and averages of all 20 frames were used for image processing. A total number of ~291,000 particles were analyzed from 6576 selected frame-averaged micrographs. Particles were picked using the ara-autopick and ara-crop tools in *Arachnid* (**Langlois** *et al.*, 2014), and the contrast transfer function parameters were estimated using the sp-defocus tool in *Arachnid* (**Langlois** *et al.*, 2014). 3D classification was performed using *RELION* software (version 1.2) (**Scheres**, 2012) to discard defective particles and identify structurally homogeneous subsets.

Initial *RELION* 3D classification, with K = 10 classes and an angular sampling of 1.8°, revealed two major populations, namely 80S ribosomes that are complete, and 80S ribosomes that lack densities for uL16 r-protein and/or eS1 r-protein, (denoted as ‘complete 80S’ and ‘incomplete 80S’, respectively). Since reconstructions from both populations showed fragmented densities, we performed a second tier of exhaustive 3D classification to further explore the heterogeneity. We quantitatively and qualitatively analyzed the classification results by (i) quantitative jumper analysis (**Chen** *et al.*, 2014) and (ii) prior structural knowledge of *S. cerevisiae* 80S ribosomes. In this way, we re-grouped classes which are conformationally and compositionally similar, and performed *RELION* auto-refinement on each class. (Figure S1).

In total, we found 10 major subpopulations, 5 complete 80S populations and 5 incomplete 80S populations. For the population of complete 80S ribosomes, we have (i) rotated 80S bound with P/E tRNA, with an average resolution of ~5.6Å, (ii) rotated 80S with P/E tRNA, ~6.3Å, (iii) rotated 80S with P/E tRNA, ~7.6Å. (vi) non-rotated 80S with P/P- and E/E- tRNAs, ~6.5Å, (v) non-rotated 80S without any tRNAs, ~6.4Å. The first three rotated 80S classes are different from each other by the degree of inter-subunit rotation and head movements, in comparison with class 4, nrt80S-P/E (Figure S1 and Table S3).

For the subpopulations of incomplete 80S ribosomes, we found (i) rotated 80S, lacking only the density for uL16, with an average resolution of ~9.4Å, (ii) rotated 80S, lacking densities for both uL16 and eS1, ~6.1Å, (iii) non-rotated 80S lacking densities for both uL16 and eS1, ~6.5Å, (vi) non-rotated 80S lacking densities for both uL16 and eS1, as well as densities for small fragments of rRNA, ~9.2Å, (v) non-rotated 80S lacking densities for both uL16 and eS1, as well as densities for small fragments of rRNA, ~9.8Å (**Fig. 5**, **Fig. S1** and **Table S4**).

Resolutions reported throughout were based on the ‘gold standard’ protocol along with the FSC = 0.143 criterion, and involved soft masking and high-resolution noise substitution (**Chen** *et al.*, 2013; **Scheres**, 2012). For the final visualization, all density maps were corrected for the effects of a soft mask in *RELION* post-processing (**Chen** *et al.*, 2013; **Scheres**, 2012), and sharpened by application of an automatically estimated negative b-factor (**Scheres**, 2012).

